# Optimization of the flash extraction of flavonoids from the leaves of *Salix babylonica* using the response surface method and an evaluation of the leaves’ high antioxidant activity

**DOI:** 10.1101/423095

**Authors:** Huijie Chen, Lei Diao, Yue Zhang, Haixin Liu, Ming Zhong, Guangxing Li

## Abstract

Many biological activities of *Salix babylonica* leaves are attributed to the plants’ high total flavonoid content. Flash extraction has the advantages of high efficiency and maximum retention of the active ingredient. In this study, flash extraction was used to extract the total flavonoids, and a Box–Behnken design was used to optimize the extraction conditions for the first time. The effects of four independent variables, including ethanol concentration, extraction voltage, time, and ratio of liquid to material on flavonoid yield, was determined, and the optimal conditions for flavonoid extraction were evaluated using response surface methodology. Statistical analyses showed that the linear and quadratic terms of these four variables had significant effects. The fitted second-order model revealed that the optimal conditions consisted of an ethanol concentration of 67.91%, extraction time of 87 s, extraction voltage of 116 V and ratio of liquid to material of 42.79. Under the optimum conditions, the experimental value of 66.40±0.80% nearly coincided with that predicted by the model. In the ferric reducing antioxidant power (FRAP) and 2,2-diphenyl-1-picrylhydrazyl radical (DPPH.) assays, the extracts showed significant antioxidant and scavenging capacity for free radicals, respectively. This study helps to better exploit the resources of *Salix babylonica* leaves and provides new insights for effective extraction of flavonoids.

## Introduction

Currently, there are increasing applications of flavonoids in food and medicine for their extensive biological functions in terms of antibacterial, antiviral and antioxidant activities [1-5]. As a biological response modifier, most of these flavonoids are derived from traditional Chinese medicine, including *Salix babylonica*. This plant is economically notable and is distributed throughout Asia, Europe, and the Americas. The leaf of *Salix babylonica* has been studied in China for antifungal and anti-obesity utilization due to its high content of flavonoids [6, 7]. Currently, there are limited studies on the extraction process and antioxidant activity of the total flavonoids from *Salix babylonica* leaves [8], and the primary focus is on the enrichment and purification of the total flavonoids. In addition, most of the extraction methods continue using the outmoded methods of reflux extraction, ultrasonic extraction and microwave extraction [9, 10]. In these cases, extraction required a long time and was labor-and material-intensive, and the most important factor was that the active ingredients of the extract were easily degraded. To use *Salix babylonica* leaf resources more optimally, there is an urgent need to establish the optimum procedure for the effective extraction of flavonoids.

The flash extraction method is a new method in recent years [11, 12]. This method uses the principle of high-speed shear force and molecular filtration to break up the stems, leaves, roots and other materials of plants into tiny particles such that the concentration of the extraction solvent reaches equilibrium in a short time [13, 14]. In addition, the solute transfer process is completed in tens of seconds or even seconds, and the active ingredients in the plant are only destroyed to a small extent. Compared with the traditional hot water extraction, enzyme extraction and other physical field-assisted extraction methods, such as microwave, ultrasonic, and ultrahigh pressure extraction, the flash extraction method uses small amounts of solvent, a short extraction time and is highly efficient [15-18]. In recent years, there have been many reports of the use of this technology to extract phytochemicals. It was used to extract water-soluble components, such as polyphenols [19], glycosides [20], and polysaccharides [21], as well as water-insoluble components, such as tannins [22], peony [23], and volatile oil [24]. Therefore, the flash extraction method with its unique advantages of speed, energy, and the savings in solvents is bound to exert greater potential for herbal extractions.

In humans, oxidation is involved in the generation of energy, but the body may be affected by a variety of factors, such as illness, aging, and free oxygen radicals produced by oxidation, that are uncontrollable and harmful to the human body. Antioxidants, such as flavonoids, can eliminate these free radicals, reduce the risk of death from related diseases, and slow aging [25, 26]. Therefore, the preparation and application of antioxidants to protect the body from free radical damage is popular [27]. In this study, flash extraction and response surface methodology (RSM) were combined to seek an effective extraction procedure. In addition, the antioxidant capacity of the extracts was investigated to verify the effectiveness of flash extraction. The results indicated that the flash extraction method could maximize the protection of the active ingredients of flavonoids with high biological function. These findings will provide a theoretical basis to develop and utilize *Salix babylonica* leaf resources.

## Materials and methods

### Materials

The leaves of *Salix babylonica* were collected in Jilin Province (China). Rutin and oligomeric proanthocyanidins (OPC) standards were purchased from the National Institutes for Food and Drug Control (Beijing, China). Aluminum nitrate, sodium hydroxide, ethanol, sodium nitrite, hexahydrate ferric chloride and ferrous sulfate were analytically pure and obtained from the Tianjin Reagent Company (Tianjin, China). 2,4,6-tri(2-pyridyl)-1,3,5-triazine (TPTZ) and 2,2-diphenyl-1-picrylhydrazyl radical (DPPH) were purchased from J&K Scientific Co. (Beijing, China).

### Extraction of total flavonoids

The leaves of *Salix babylonica* were crushed using a grinder and passed through a 20-mesh screen. With the use of a flash extractor, a number of samples were accurately weighed and extracted under different extraction times, extraction numbers, ratio of liquid to raw material, ethanol concentration and extraction voltage. The extract was separated from the insoluble residue using centrifugation (3000 rpm, 5 min), and the extract was filtered using a 0.45 μm microporous membrane. The filtrate was concentrated and dissolved to 100 mL to determine the concentration of the flavonoids.

### Determination of the flavonoid contents

The colorimetric method was used to determine the content of the flavonoids in the extract [28, 29]. One milliliter of flavonoid extract, 1 mL of 5% (w/w) sodium nitrite and 5 mL of 80% (v/v) ethanol were accurately measured and mixed for 6 min. One milliliter of 10% (w/w) aluminum trichloride were added and incubated for 6 min, and 10 mL of 1 mol/L sodium hydroxide was added. The absorbance of the solution was measured using a UV-2900 spectrophotometer at 510 nm after 15 min. A blank control was used. The standard curve (*y* = 0.0164 + 0.0097*x*, where *y* is absorbance value of sample, and *x* is sample concentration) ranged from 16.32–65.28 μg/mL (*R*^*2*^=0.9993).

### Experimental design

The extraction rate of the flavonoids is influenced by many factors, such as the extraction voltage, extraction time, the ratio of liquid to raw material, ethanol concentration and extraction number [30, 31]. Based on the single factor test results, factors that have a significant impact on the flavonoids were selected to test using response surface methodology. The four significant impact factors were designated *X*_*1*_, *X*_*2*_, *X*_*3*_, and *X*_*4*_, and each factor was set at three levels, coded +1, 0, −1 for high, medium and low values, respectively (Table 1). Four variables were encoded based on the following equation:

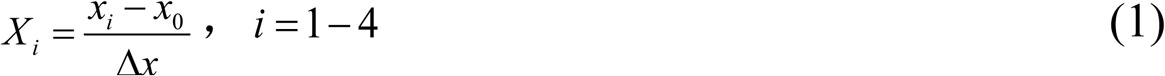

where *X*_*i*_ represented the encoded value of the variable; *x*_*i*_ represented the actual value of the variable; *x*_*0*_ represented the actual value of the independent variable at the center point, and Δ*x* represented the change value of the variable, respectively.

**Table 1.** Variables and design levels of response surface methodology.

| Independent variables | Coded symbols | Levels |  |  |
| --- | --- | --- | --- | --- |
|  |  | -1 | 0 | 1 |
| Extraction voltage (V) | $X_1$ | 110 | 120 | 130 |
| Ethanol concentration (%) | $X_2$ | 50 | 60 | 70 |
| Extraction time (s) | $X_3$ | 80 | 90 | 100 |
| Ratio of liquid to material | $X_4$ | 30 | 40 | 50 |

To use the Box-Behnken design and obtain the best extraction conditions, a second order polynomial model was used to illustrate the relation between the response values and independent variables. The model is as follows:

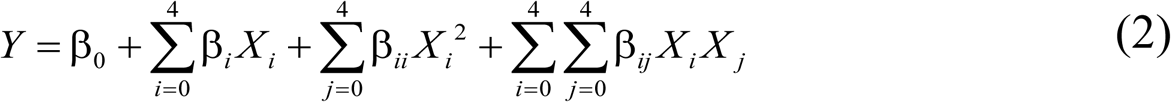

Where *Y* represented the response function; *β*_*0*_ represented the constant, and *β*_*i*_, *β*_*ii*_ and *β*_*ij*_ represented the coefficients of the linear, quadratic and interactive terms, respectively. *X*_*i*_, X_i_^2^ and*X*_*i*_*X*_*j*_ were the encoded independent variables, interaction and quadratic terms, respectively.

To design the experiments and analyze the experimental data, design-expert software (version 8.0) was used. Based on the analysis of variance, the regression coefficients of individual linear, quadratic, and interaction terms were determined. The regression coefficients were used to provide statistical calculations to generate dimensional and contour maps from the regression model. The lack of fit and the coefficient of determination (*R*^*2*^) were generated using the Design-Expert software. They represented the accuracy of the model. Fischer’s F-test at a probability (*P*) of 0.001, 0.01 or 0.05 examined the significance of the model, as well as the encoded independent variables, interaction and quadratic terms.

### Antioxidant activity assay of flavonoids

A ferric reducing antioxidant power (FRAP) experiment was used to measure the total antioxidant activity of the extracts and standards (FeSO_4_ solution). A 2,2-diphenyl-1-picrylhydrazyl radical (DPPH) experiment was used to measure the free radical scavenging activity of the extracts and standards using OPC.

### FRAP assay

FRAP reagents included 50 mmol/L acetate buffer, which contained 20.4 g C_2_H_3_NaO_2_ and 80 mL C_2_H_4_O_2_ per liter, 10 mmol/L TPTZ (2,4,6-tripyridyl-s-triazine) solution in which 40 mmol/L HCl solution was used as the solvent, and 20 mmol/L FeCl_3_·6H_2_O. The working FRAP reagent was obtained by mixing 100 mL acetate buffer, 10 mL TPTZ solution, and 10 mL FeCl_3_·6H_2_O solution [32]. One hundred microliters of different amounts of FeSO_4_ solutions or extracts were added to 6 mL FRAP reagent and incubated in a 37^°^C water bath for 30 min. The wavelength of 593 nm was used to monitor the absorbance of the samples.

### DPPH assay

Sixty micromolar DPPH was dissolved in 3 mL ethanol, and 0.5 mL of different amounts of extracts or OPC were added. Blank experiments, which contained 0.5 mL of 99% ethanol instead of the extract, were also conducted. The absorbance values were monitored at 517 nm [33]. The inhibition rate (IR) of the DPPH free radical was calculated using the following equation [34]:

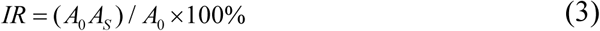

where *A*_*0*_ was the absorbance of the blank experiments, and *As* was the absorbance in the presence of the samples. When the IR is 50%, the corresponding concentration is called the inhibition concentration 50 (IC_50_). Therefore, the value of the IC_50_ was obtained by fitting the sample concentration and inhibition rate [35].

### Statistical analysis

Statistical analysis of the single-factor experimental data was performed using SPSS 11.5 software (SPSS Inc., Chicago, IL, USA). Dunnett’s one-tailed t-tests permitted the examination of the statistical significance of the means in the different levels of parameters. Stat-Ease Design-Expert 8.0.0 (Trial version, Stat-Ease Inc., Minneanopolis, MN, USA) was used for the experimental design and regression analysis of the experimental data. Student’s t-test permitted the determination of the statistical significance of the regression coefficient, and Fischer’s F-test determined the second-order model equation at a probability (*P*) of 0.001, 0.01 or 0.05. The adequacy of the model was determined by evaluating the lack of fit, the coefficient of determination (R^2^), and the F-test value obtained from the analysis of variance (ANOVA) that was generated. All the experiments were conducted in triplicate, and the average values ± SD (standard deviation) were calculated.

## Results

### Single factor test results

### Effect of extraction voltage in single factor analysis

The extraction voltage was one of the important factors affecting the yield of total flavonoids [36]. The effects of different voltages (60, 80, 100, 120, and 140 V) on the yield of total flavonoids were investigated, while the other extraction conditions were as follows: ethanol concentration of 70%, ratio of ethanol to raw material of 30:1, extraction time of 60 s, and extraction number of 3. The results showed that the yield increased greatly when the voltage increased from 60 to 140 V, and the highest yield was obtained at 120 V (Fig 1A). With the increase in voltage, the yield of flavonoids increased. This phenomenon might be attributable to the increase of the rotational speed of the head of the flash extractor, the increase in the temperature of the extraction solvent at the same time, and the flavonoids undergoing side reactions, such as oxidation or decomposition, resulting in a decrease in the content of flavonoids [36]. In this experiment, the optimal extraction voltage was considered to be 120 V.

**Fig 1.**
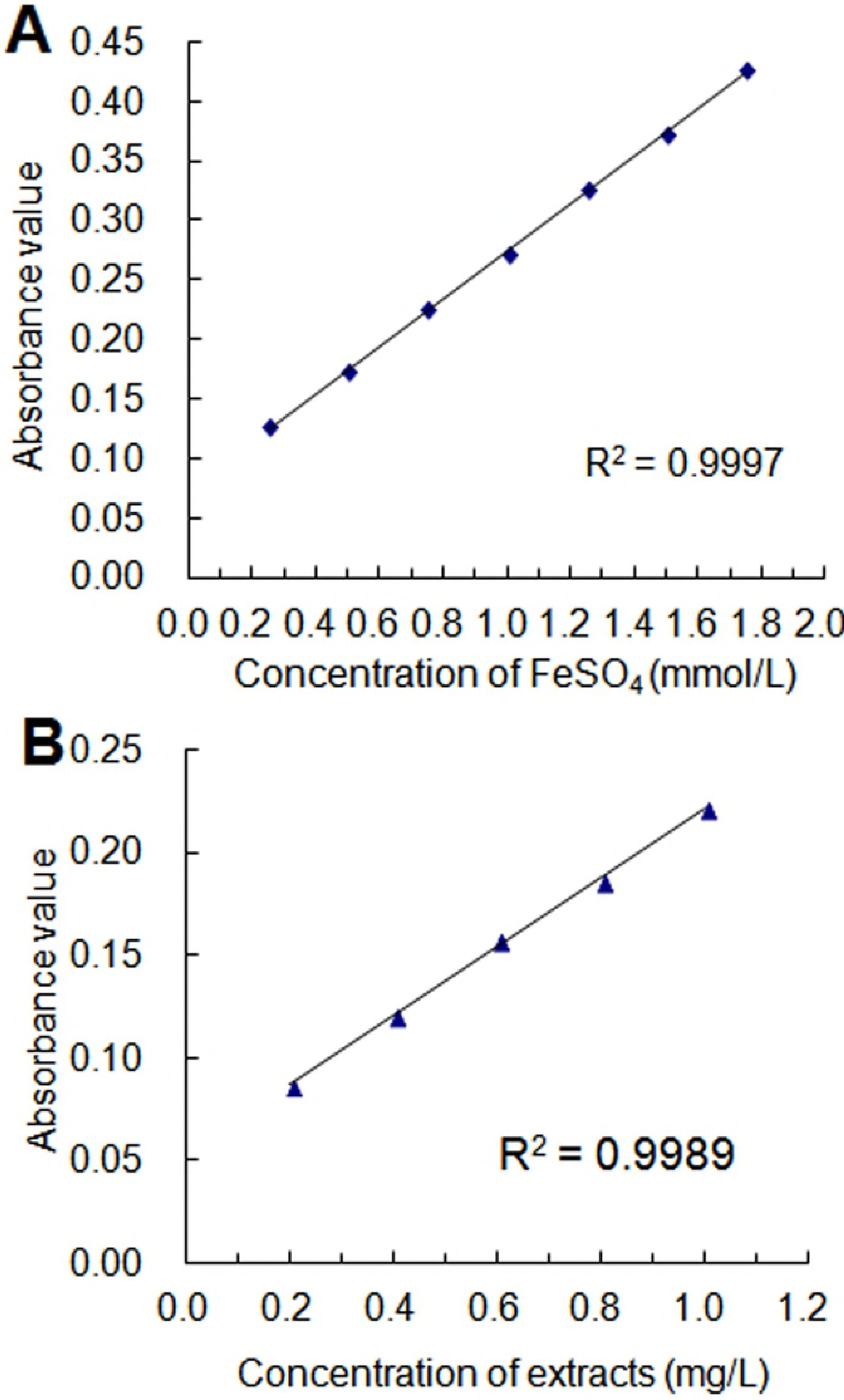
Column chart showing the effect of different factors on the yield of total flavonoids. The effect of (A) different voltage, (B) ethanol concentration, (C) ratio of ethanol to raw material, (D) time and (E) extraction number on the yield of flavonoids was determined. (t-test, *\* p* < 0.05, *\*\* p* < 0.01, *\*\*\* p* < 0.001)

### Effect of ethanol concentration in single factor analysis

Different concentrations of ethanol (40, 50, 60, 70, and 80%) were utilized to study the effect of the ethanol concentration on the extraction rate, while the other experimental parameters were as follows: the extraction voltage was 120 V, the ratio of liquid to material was 30, the extraction time was 60 s, and the number of extraction was 3.

As shown in Fig 1B, the extraction yield of the flavonoids was greatly influenced by the concentration of ethanol [37, 38]. This yield increased with the augmentation of the ethanol concentration until it reached a maximum production of 37.53 ± 1.12 mg/g at 60% of ethanol. It also indicated that the maximum extraction yield was obtained with 60% ethanol. Therefore, 60% ethanol was used as the center point for the additional RSM experiments.

### Effect of the ratio of liquid to material in single factor analysis

The effect of the ratio of material to liquid on the yield of flavonoids is also important [39, 40]. Different ratios of liquid to material (20, 30, 40, 50 and 60) were designed to investigate their effect on the extraction rate. Simultaneously, the other conditions were as follows: extraction voltage of 120 V, ethanol concentration of 60%, extraction time of 60 s, and number of extraction of 3. As displayed in Fig 1C, the yield increased strongly as the ratio increased from 20 to 60. When the ratio of the liquid to the material continued increasing, the output was maintained at a stable level. Therefore, the ratio of liquid to material of 40 was used in this study.

### Effect of extraction time in single factor analysis

Using a fixed extraction voltage (120 V), ethanol concentration (60%), ratio of liquid to material (40) and extraction number (3), the effects of different extraction times (30, 60, 90, 120, and 150 s) on the extraction rate of the flavonoids were determined. The yield of the flavonoids increased significantly with the increase in extraction time from 30 s to 150 s (Fig 1D). However, at more than 90 s, the flavonoid production decreased slightly. This finding may be attributable to the prolonged extraction time; a large amount of heat is generated due to the rapid rotation of the head of the flash extractor; the temperature of the extraction solvent is increased, and the flavonoids undergo side reactions, such as oxidation or decomposition, resulting in a decrease in the content of the flavonoids [41]. Therefore, 90 s was used as the center of the extraction time for RSM.

### Effect of the extraction number in single factor analysis

The number of the extraction was also important to extract the active compound [42, 43]. The number of the extraction was related to the extraction efficiency, cost, and flavonoid yield. Respectively, different extraction numbers (1, 2, 3, 4 and 5) were used to extract the flavonoid under the optimal parameters obtained above. As shown in Fig 1E, with the increase in the number of the extraction from 1 to 2, the yield of flavonoids increased significantly. However, when the number of extractions was more than 2, the increase in flavonoid production was not significant. Considering the efficiency of extraction, the number of extraction was set at 2 in this study.

### Model fitting and statistical analysis of the extraction process optimization

Response surface methodology optimization is more advantageous than the traditional single parameter optimization, because it saves time, space and raw material. To optimize the four individual parameters of the Box-Behnken design, the 27 experiments designed were completed. Using multiple regression analyses, the following second order polynomial equations are used to correlate the response variables and test variables:

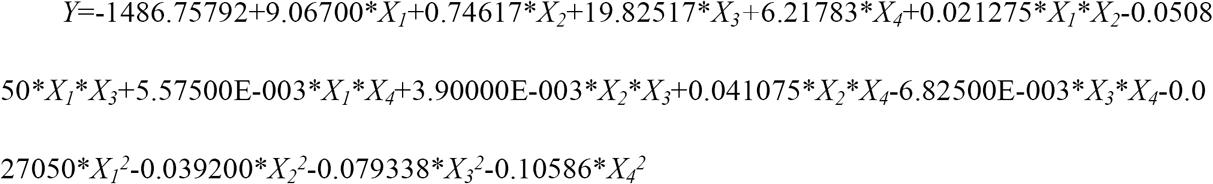

where *X*_*1*_, *X*_*2*_, *X*_*3*_ and *X*_*4*_ were the encoded values on extraction voltage, ethanol concentration, extraction time and the ratio of liquid to material, respectively.

To determine whether the model was significant, the significance of each coefficient was determined using the *F*-test and *p*-value in Table 2. Smaller *p*-values indicated higher significances for the corresponding coefficient [44]. In this study, the ANOVA of the quadratic regression model demonstrated that the model was highly significant, as was evident from the *F*-test with a very low probability value (*p* < 0.0001). With the determination coefficient (*R*^*2*^ = 0.9755) close to 1, this finding indicated that the actual values had a strong correlation with the predicted values [45]. With Adj. *R*^*2*^ values of 0.9469, it could be concluded that more than 90% of the total flavonoid content could be predicted. Only 5% could not be predicted by the model.

**Table 2.** ANOVA table for the response surface quadratic model.

| Source | Sum of squares | Degree of freedom | Mean square | F-Value | $p$-Value (Prob > F) | Significant |
| --- | --- | --- | --- | --- | --- | --- |
| **Model** | 2163.56 | 14 | 154.54 | 34.10 | <0.0001 | *** |
| $X_1$ | 302.40 | 1 | 302.40 | 66.73 | <0.0001 | *** |
| $X_2$ | 416.54 | 1 | 416.54 | 91.91 | <0.0001 | *** |
| $X_3$ | 427.09 | 1 | 427.09 | 94.24 | <0.0001 | *** |
| $X_4$ | 86.24 | 1 | 86.24 | 19.03 | 0.0009 | *** |
| $X_1X_2$ | 18.11 | 1 | 18.11 | 4.00 | 0.0688 | |
| $X_1X_3$ | 103.43 | 1 | 103.43 | 22.82 | 0.0005 | *** |
| $X_1X_4$ | 1.24 | 1 | 1.24 | 0.27 | 0.6100 | |
| $X_2X_3$ | 0.61 | 1 | 0.61 | 0.13 | 0.7204 | |
| $X_2X_4$ | 67.49 | 1 | 67.49 | 14.89 | 0.0023 | ** |
| $X_3X_4$ | 1.86 | 1 | 1.86 | 0.41 | 0.5334 | |
| $X_1^2$ | 39.02 | 1 | 39.02 | 8.61 | 0.0125 | * |
| $X_2^2$ | 81.95 | 1 | 81.95 | 18.08 | 0.0011 | ** |
| $X_3^2$ | 335.70 | 1 | 335.70 | 74.08 | <0.0001 | *** |
| $X_4^2$ | 597.70 | 1 | 597.70 | 131.89 | <0.0001 | *** |
| **Residual** | 54.38 | 12 | 4.53 |  |  |  |
| **Lack of fit** | 27.50 | 10 | 2.75 | **0.20** | **0.9760** | **Not significant** |
| **Pure error** | 26.89 | 2 | 13.44 |  |  |  |
| **Cor total** | 2217.94 | 26 |  |  |  |  |
| $R^2$ | **0.9755** | | | | | |
| **Adj. $R^2$** | **0.9469** |  |  |  |  |  |
| **Pred. $R^2$** | **0.9013** |  |  |  |  |  |
| **Adequate**<br>**precision** | **18.194** |  |  |  |  |  |

(t-test, \* $p < 0.05$ , \*\* $p < 0.01$ , \*\*\* $p < 0.001$ )

The failure of the model was represented by the lack-of-fit shown in Table 2. The lack of fit of the *F*-value was 0.2000, and the *P*-value was 0.9670, which illustrated that the pure error was not significantly relative. In summary, the model was appropriate. Simultaneously, the adequate precision of 18.194 indicated that the model discrimination was adequate. Therefore, this study model could be used to navigate the design response surface space.

The *P*-values of each model term presented in Table 2 indicated that the total flavonoid yield was significantly affected by four independent variables (*X*_*1*_, *X*_*2*_, *X*_*3*_, *X*_*4*_) and all of the quadratic terms (*X*_*1*_^*2*^, *X*_*2*_^*2*^, *X*_*3*_^*2*^, *X*_*4*_^*2*^). In addition, the ethanol concentration and extraction time were the most significant factors on the total flavonoid yield followed by the extraction voltage and the ratio of liquid to material.

### Response surface analysis of extraction process optimization

The interaction effects of the factors in response surface experiments were displayed using three-dimensional response surface plots (Fig 2) and two-dimensional contour plots (Fig 3). The former described the sensitivity of the variable change on the response value, and the latter illustrated the important coefficients between the different variables [46, 47]. The impact of the two factors on the response could be displayed at one time by these plots. In these figures, the other two factors remained at level zero.

**Fig 2.**
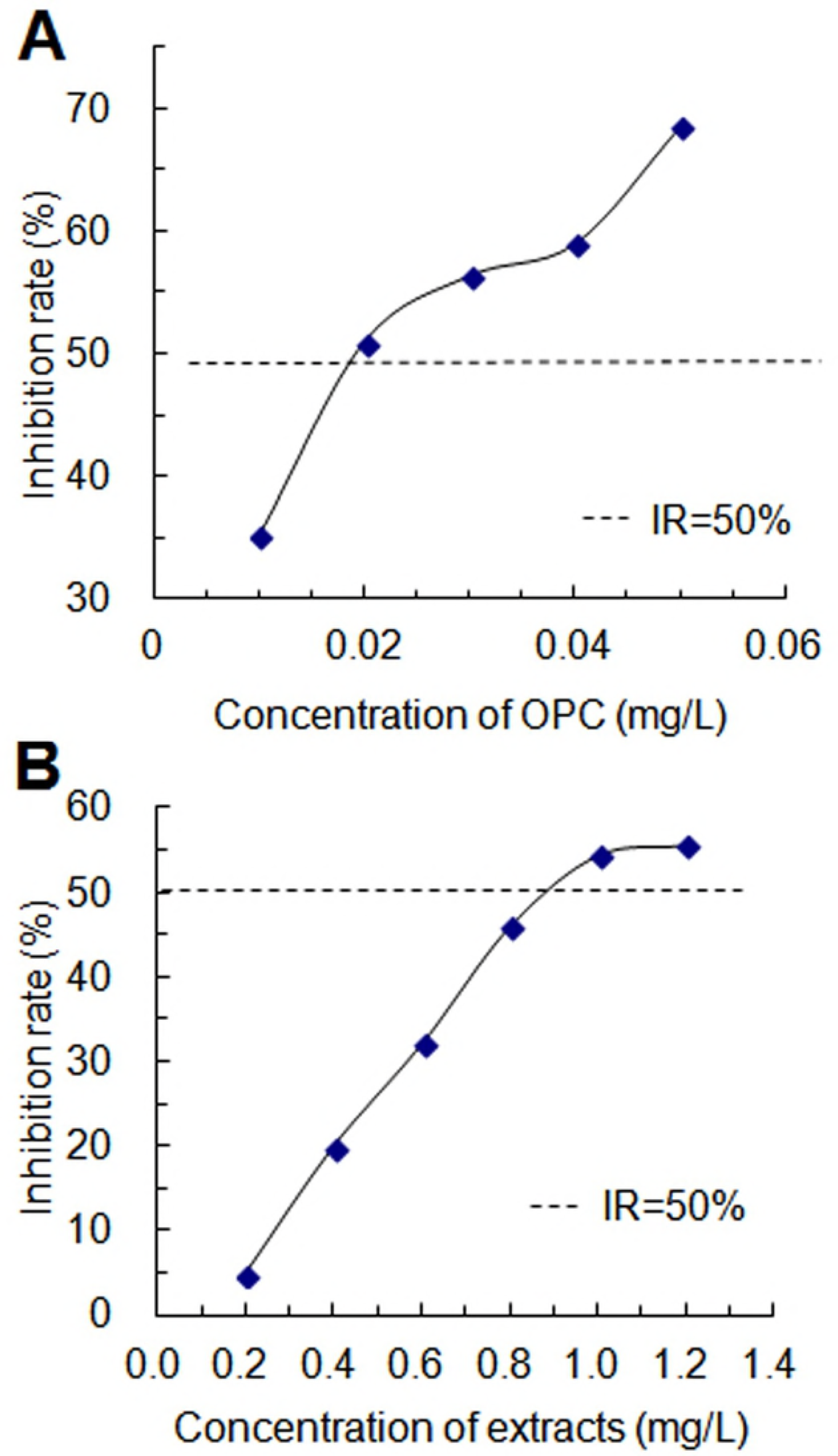
Response surface method analysis (3D) of the different interaction factors.

**Fig 3.**
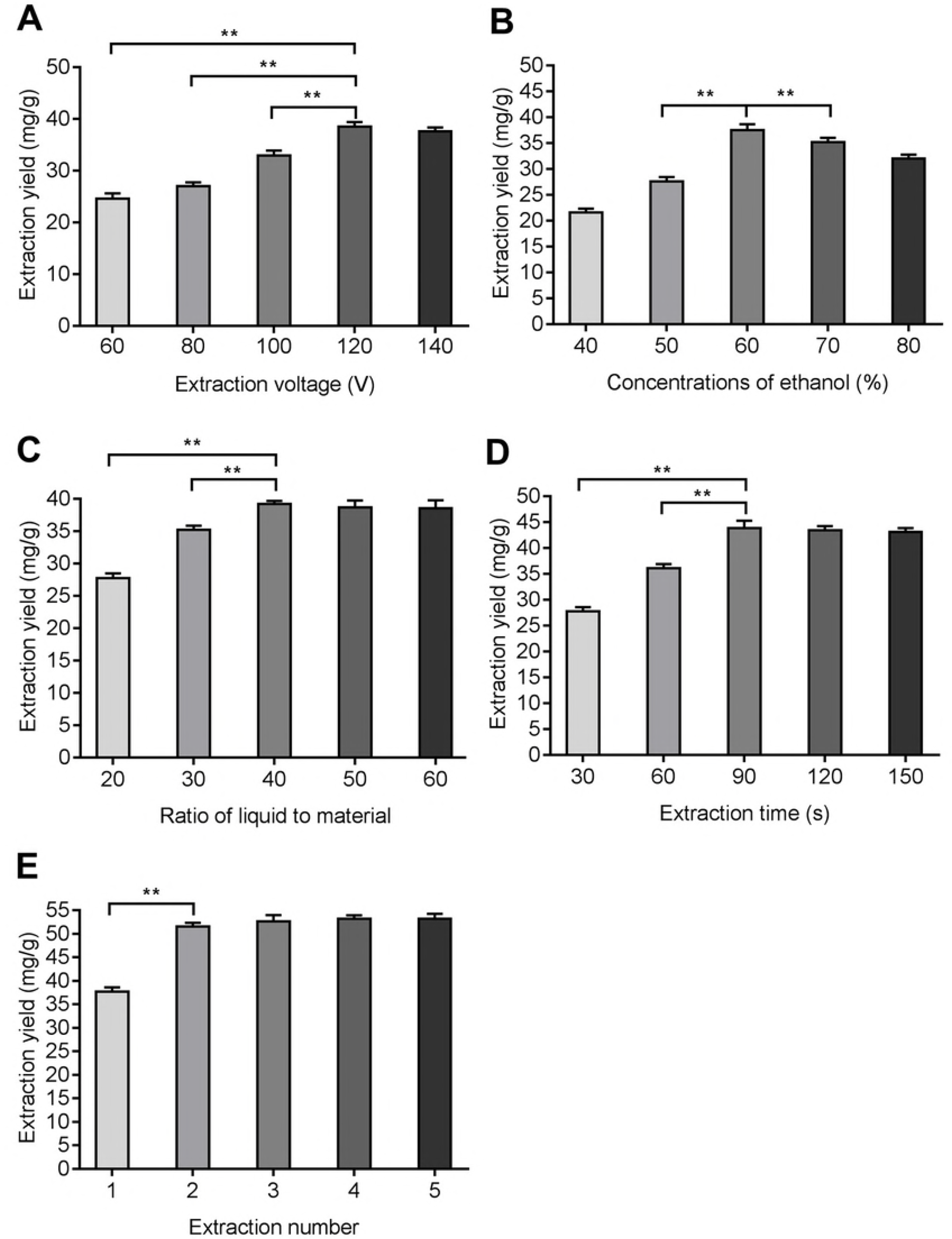
Contour plots (2D) of the different interaction factors.

The influences of the extraction voltage and ethanol concentration on the flavonoid yield are shown in Figs 2A and 3A. When the ethanol concentration was less than 68%, the flavonoid yield increased, while when it was higher than 68%, the flavonoid yield decreased, and when the voltage was low, the yield changed more significantly. When the ethanol concentration increased from 50% to 60%, the extraction voltage had little effect. However, when the ethanol concentration increased from 65% to 70%, the voltage had a greater effect on the flavonoid yield.

Figs 2B and 3B showed the reciprocal interactions of the extraction voltage and extraction time on the yield of flavonoids. The mutual interactions were significant. With the increase of extraction voltage, the flavonoid yield clearly decreased, especially when the extraction time was longer. After a period of time, the flavonoid yield increased. However, at more than this range, the output did not increase noticeably. Simultaneously, when the extraction voltage was raised to 116 V, the extraction time had a significant effect on the flavonoid yield.

The effect of extraction voltage and ratio of liquid to material on the flavonoid yield is shown in Figs 2C and 3C. With the extraction voltage dropping, when the ratio of liquid to material was relatively low, there was little effect on the flavonoid yield. However, when the ratio of liquid to material was relatively high, the decrease in voltage had a greater effect on the flavonoid yield. When the ratio of liquid to material increased from 30 to 42.79, the flavonoid yield increased significantly to 42.79 and began to decline. Therefore, in order to access the maximum increase, the ratio of liquid to material was more effective at 42.79.

The influences of ethanol concentration and extraction time on the yield of flavonoids are visible from Figs 2D and 3D. This result showed that the extraction time exhibited a significant effect on the flavonoid yield, while the ethanol concentration had a weaker effect. With the extraction time increasing to 87 s, the flavonoid extraction rate increased and then decreased. When the ethanol concentration increased from 50% to 67.79%, the extraction of the flavonoids increased significantly, while at more than 67.79% it decreased. Therefore, to achieve maximum increase, the ethanol concentration used was 67.79%.

The effects of the mutual interactions between the ethanol concentration and the ratio of liquid to material were significant (Figs 2E and 3E). With the ethanol concentration increasing from 50% to 67.79%, the flavonoid production increased linearly, but at more than 67.79%, it decreased. This finding was probably attributable to the scope of the solvent; the polarity was appropriate and more suitable to extract flavonoids [48]. When the ratio of liquid to material increased from 30 to 42.79, the flavonoid yield increased at first and then decreased.

In Figs 2F and 3F, the impact of the interaction of the ratio of liquid to material and extraction time was displayed. An appropriate extraction time was essential for the extraction of flavonoids. When the extraction time increased from 80 to 87 s, the flavonoid yield increased, but when the extraction time was longer than 87 s, the yield decreased. The ratio of liquid to material was lower, and the yield also stayed lower. When the ratio of the liquid to material increased from 30 to 42.79, the flavonoid yield increased significantly, and after 42.79, the change was very small.

### Validation of the model

As shown in Table 3, the optimum extraction conditions were obtained, and the flavonoid yield predicted by the Design Expert Software was 66.54 mg/g. To validate the adequacy of the model equations, a verification experiment was conducted under the optimal conditions (within the experimental range): extraction voltage 116 V, ethanol concentration 68%, extraction time 87 s, the ratio of liquid to material 43 and extraction number 2, respectively. A mean value of 66.40±0.80 mg/g (N=3) was obtained from the practical experiments. The average was close to the predicted value of the RSM model. Therefore, this model could be used to extract flavonoids, and the yield of flavonoids was higher than that of other extraction conditions.

**Table 3.** Experimental and predicted values of the model under optimal conditions.

| Optimum<br>condition |  |  |  | Extraction yield |  |
| --- | --- | --- | --- | --- | --- |
|  |  |  |  | (mg/g) |  |
| Extraction<br>voltage (V) | Concentration<br>of ethanol (%) | Extraction<br>time (s) | Ratio of liquid<br>to material | Experimental | Predicted |
| 116 | 67.91 | 87.00 | 42.79 | **$66.40 \pm 0.80$** | **66.54** |

### FRAP assay

As shown in Fig 4, a linear correlation between the concentration of the FeSO_4_ solution or extracts and the absorbance was displayed. In addition, the concentration of extracts was expressed as milligrams of *Salix babylonica* leaves per liter. In addition, the total antioxidant activity of the 1 mg extracts approximately equaled the total antioxidant activity of 0.5–0.75 mmol FeSO4.

**Fig 4.**
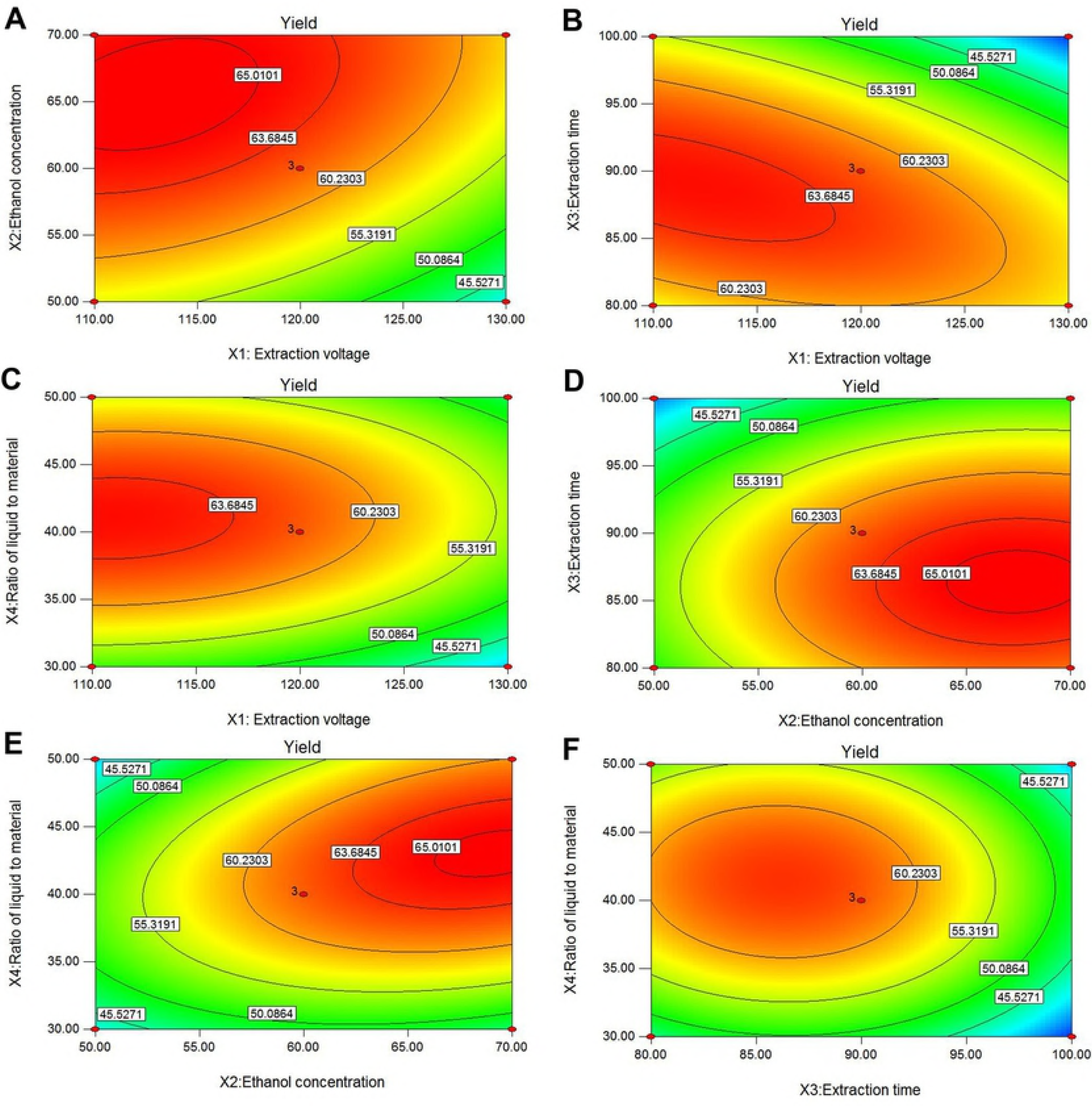
Total antioxidant activity of (A) FeSO_4_ solutions and (B) extracts.

### DPPH assay

DPPH is a stable and well-characterized solid radical source, and it is a traditional and the most popular free radical used to assay free radical scavenging activity [49]. OPC, which has a strong ability to scavenge a free radical, was used as the standard to evaluate the free radical scavenging activity of the extracts. As shown in Fig 5, the inhibition rate (IR) and IC_50_ were used to characterize the free radical scavenging activity of the extracts. With the concentration increasing, the value of the IR increased. The IC_50_ value of the extracts from the *Salix babylonica* leaves was 0.8937 mg/L, while the IC_50_ value of the OPC was 0.0196 mg/L. Therefore, the free radical scavenging activity of the 1 mg extracts was approximately equivalent to that of 0.021 mg OPC.

**Fig 5.**
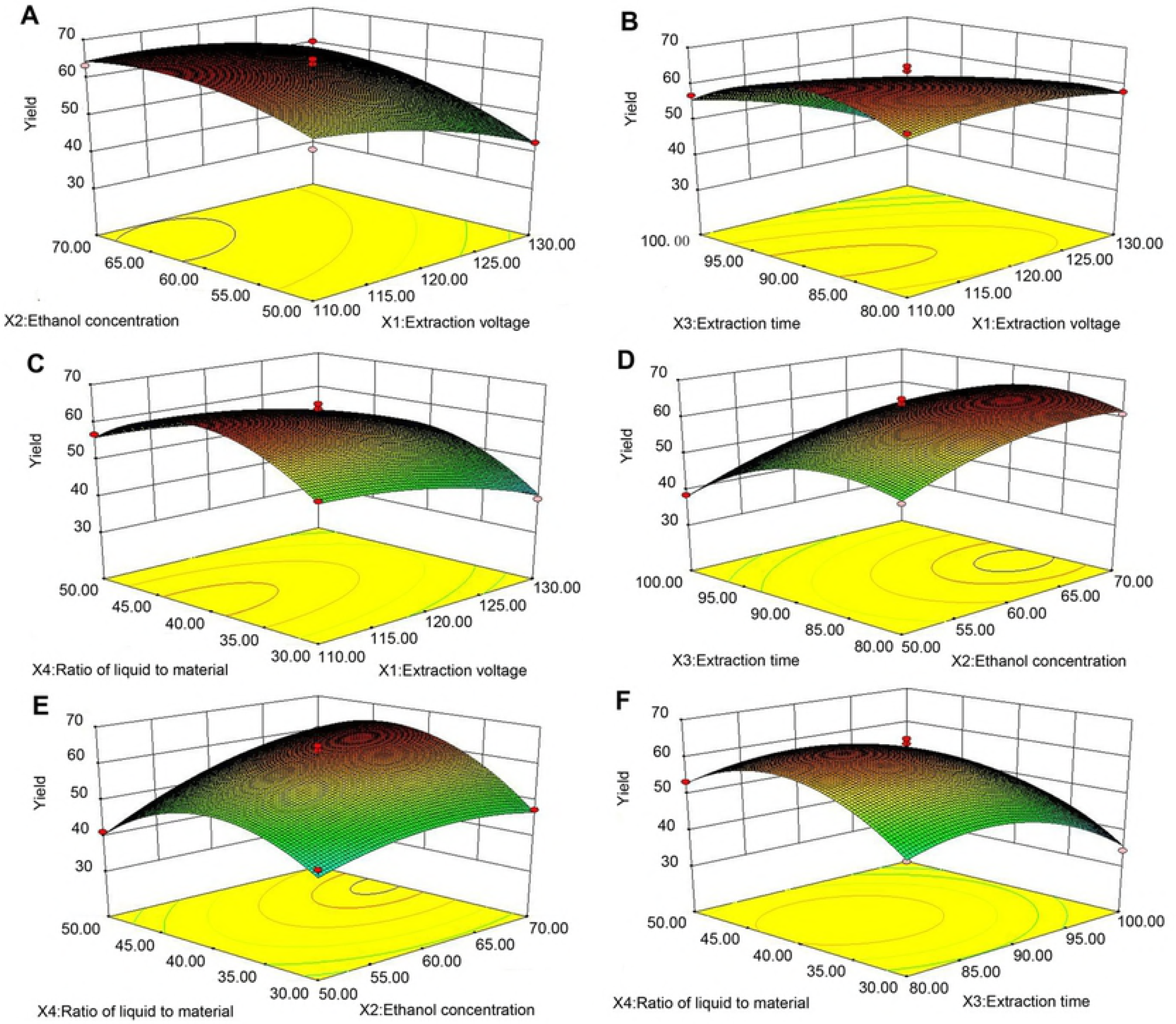
Scavenging activity of (A) OPC and (B) extracts on DPPH free radicals.

## Discussion

Belonging to *Salicaceae* and *Salix*, *Salix babylonica* is a fast-growing deciduous tree, which is widely cultivated in Asia, Europe and America. In foreign countries, the leaves of these trees are an important source of feed for ruminants, particularly in areas that experience harsh environmental conditions. Plant extracts from *Salix babylonica* leaves contain saponins and other secondary natural metabolites, which can improve nutrient digestibility and feed utilization [50-53]. In China, its leaves, branches, and floes are all included in the Chinese Dictionary of Traditional Chinese Medicine as a traditional Chinese medicine. The Compendium of Materia Medica recorded the following: *Salix babylonica* is the lower grade of this product. This plant is cold in nature and tastes bitter. The plant has the functions of hurricane, diuretic, analgesic, and swelling. According to reports, *Salix babylonica* is rich in flavonoids [54, 55], which have diverse physiological activities, such as anti-inflammatory, anti-tumor, antioxidant, anti-dementia and anti-obesity [56-64]. It must have widespread utilization in the medicine and food industry in the future.

*Salix babylonica* is rich in resources and cultivated throughout China, but its medicinal value has not been properly developed and utilized to date. The optimal extraction procedures of the flavonoids from *Salix babylonica* leaves with new techniques had not been studied frequently, and most of the extraction methods are limited to conventional extraction methods with low efficiency and obvious shortcomings. Xiaoyan Yi *et al.* used the reflux extraction method, and the yield of flavonoids was 5.15% [10]. Keyue Liu *et al.* adopted the microwave extraction method, and the flavone yield was 5.67% [65]. The flash extraction method, also known as the tissue fragmentation extraction method, is an emerging extraction method [66-68]. The flash extraction method is applicable to the extraction of the roots, stems, leaves, flowers, fruits, seeds (except for fine seeds) and whole plants and can be entered into the extraction system only by appropriate treatment. In addition to ether and other volatile solvents, water, methanol, ethanol, acetone, and microemulsion [69] can be directly selected as the solvent according to the desired effective site or chemical composition. The extraction process flow is simple, rapid, energy saving, and environmentally friendly. It can be implemented at room temperature, avoiding heat interference, effectively protecting the heat-sensitive components [22, 24] and has obvious advantages compared with traditional extraction methods. In traditional Chinese Medicine research, the flash extraction method will inevitably exert greater potential. In this study, the flesh extraction method was used to extract flavonoids from *Salix babylonica*. The extraction rate of the flavonoids was 60.4 mg/g (6.04%), which is significantly higher than those of the other extraction methods.

In the human body, free radicals and oxidants are produced from either normal cell metabolism or external sources. Excessive free radicals and oxidants that accumulate in the human body could generate a phenomenon called oxidative stress and result in oxidative damage to cause the deterioration of the normal function of proteins, nucleic acids and many other biomasses [70]. The antioxidant activity of the flavonoid extract of *Salix babylonica* leaf was examined by Yaoyao Su *et al*. who used the ultrasound and microwave methods to extract α-glucosidase inhibitors from weeping willow leaves and found that the inhibition rate of α-glucosidase was 77.85% [9]. Ruiping Ji *et al*. used indirect iodine quantitative methods to determine the peroxidation of animal and plant oils and fats and showed that the highest antioxidant value of *Salix babylonica* leaf was 0.204% [8]. In addition, Jie Xie *et al*. used a homogenate extraction technique to extract bamboo leaf flavonoids. The yield of total flavonoids extracted from the bamboo leaves was 8.50 mg/g, which was 20.2% higher than that of the ethanol reflux extraction. The IC50 of the total flavonoids obtained by flash extraction was 67.35 μg/mL, which scavenged free radicals. The effect is better than ethanol refluxing [71]. However, there was no report on the activity of the total flavonoids scavenging free radicals in *Salix babylonica* leaves. For the first time, this study describes the application of the flash extraction and response surface optimization methods. On the basis of improving the yield of flavonoids, the free radical scavenging activity was maintained at a high level with an IC_50_ of 0.8937 mg/L.

In this study, response surface methodology was used to establish the optimal conditions for the extraction of total flavonoids from *Salix babylonica* leaves using flash extraction. The yield of the flavonoids was maximized and increased significantly compared with those of the conventional extraction methods. In addition, antioxidant assays showed that the extracts with the established procedures in this experiment had stronger antioxidant ability, and 1 mg extracts equaled approximately 0.5–0.75 mmol FeSO_4_ or 0.021 mg OPC, respectively. These results will help to better exploit the resources of *Salix babylonica* leaves and provide new insights for the effective extraction of flavonoids with greater bioactivity.

## Acknowledgments

This research was supported by the National Natural Science Foundation of China (31172295 and 31272569). Simultaneously, this research was supported by the University Students’ Scientific and Technological Innovation Research Projects (2014088) operated by the Jilin Agricultural Science and Technology Institute of Jilin Province.

## Authors’ Contributions

**Conceptualization:** Yue Zhang, Guangxing Li.

**Formal analysis:** Yue Zhang.

**Funding acquisition:** Huijie Chen, Lei Diao, Haixin Liu, Guangxing Li.

**Investigation:** Huijie Chen, Lei Diao, Haixin Liu, Ming Zhong.

**Supervision:** Guangxing Li.

**Writing – original draft:** Huijie Chen.

**Writing – review and editing:** Huijie Chen, Lei Diao, Guangxing Li.

## References

1 Dhiman A, Nanda A, Ahmad S. A quest for staunch effects of flavonoids: Utopian protection against hepatic ailments. Arabian Journal of Chemistry. 2016; 9: S1813–S1823. https://doi.org/10.1016/j.arabjc.2012.05.001

2 Jing S, Wang SS, Li Q, Zheng L, Yue L, Fan SL, et al. Dynamic high pressure microfluidization-assisted extraction and bioactivities of Cyperus esculentus (C. esculentus L.) leaves flavonoids. Food Chemistry. 2016; 192: 319–327. https://doi.org/10.1016/j.foodchem.2015.06.097 PMID: 26304354

3 Lani R, Hassandarvish P, Shu MH, Phoon WH, Chu JJH, Higgs S, et al. Antiviral activity of selected flavonoids against chikungunya virus. Antiviral research. 2016; 133: 50–61. https://doi.org/10.1016/j.antiviral.2016.07.009 PMID: 27460167

4 Sellem I, Kaaniche F, Mtibaa AC, Mellouli L. Anti-oxidant, antimicrobial and anti-acetylcholinesterase activities of organic extracts from aerial parts of three Tunisian plants and correlation with polyphenols and flavonoids contents. Bangladesh Journal of pharmacology. 2016; 11: 531–544. https://doi.org/10.3329/bjp.v11i2.26001

5 Zhan G, Pan LQ, Tu K, Jiao S. Antitumor, antioxidant, and nitrite scavenging effects of Chinese water chestnut peel flavonoids. Journal of food science. 2016; 81(10): 2578–2586. https://doi.org/10.1111/1750-3841.13434 PMID: 27603811

6 Liu KY, Liu HJ, Zhou B, Ha LK. Studies on chemical constituents from salix babylonica L.and their stimulating lipolysis activity. Journal of Fudan University(Natural Science). 2008; 47(4): 520–523.

7 Sun GL, Yan XY. Advances in antifungal research of traditional chinese medicine. Heilongjiang medicine journal. 2015; 28(2): 295–297.

8 Ji RP, Hu BF, Han X. Preliminary study on antioxidant activity and synergistic effect of *Salix babylonica L*. leaf extract. Journal of Jinzhong university. 2014; 31(3), 52–55.

9 Su YY, Tong QY. Ultrasonic-microwave synergistic extraction of α-glucosidase inhibitor from leaves of *Salix babylonica L.* Science and technology of food industry. 2014; 35(2): 235–241.

10 Yi XY. Study on extraction process of total flavonoids from *Salix babylonica L.* leaves. Heilongjiang animal science and veterinary medicine. 2015; 9: 203–205.

11 Tao T, He F, Ji XM, Lai M, Wang PZ, Mei YN. Response surface methodology for optimization of flash extractionfor fenugreek (*Trigonella foenum-graecum L.*) polysaccharides and research of its humectant properties. Fine chemicals. 2016; 33(6): 666–673.

12 Wu DM. Application of flash extractor in traditional Chinese medicine research. Chinese Journal of Experimental Traditional Medical Formulae. 2006; 12(7): 34, 37.

13 Liu YZ, Yuan K, Ji CR. New method of extraction on the chemical components of Chinese medicinal plants-extracting method by smashing of plant tissue(EMS). Henan Science. 1993; 11(4):265–268.

14 Liu YZ. Principle and practice of smashing tissue extraction and herbal blitzkrieg extractor. Chinese journal of natural medicines. 2007; 5(6): 401–407.

15 Li JY, Liu YZ. Progress of smashing tissue extraction in research of Chinese materia medica. Chinese Traditional and Herbal Drugs. 2011; 42(10): 2145–2149.

16 Zhao XE, Geng YL, Wang DJ, Li SB, Liu JH, Wang X. Determination of Chlorogenic Acid and Luteolin in Honeysuckle by HPLC-DAD. Chinese Journal of Analysis Laboratory. 2010; 29: 77–80.

17 Liu ZY, Liu YZ, Liu GF, Zhao YQ. Study on Flash Extraction and Purification of *Gynostemma Pentaphyllum*. Chinese Traditional and Herbal Drugs. 2009; 40(7): 1071–1073.

18 Chen F, Sheng LQ, Ma JL. Smashing tissue extraction of procyanidine from pine needles and its antioxidant activity. Chinese Journal of Pharmaceuticals. 2014; 45(2): 120–123.

19 Zhu XY, Yang JH, Xie J, Dai CW, Wang MD, Wang P. Study on the technology of flash extracting tea polyphenols from green tea. Jiangsu Agricultural Sciences. 2010; 5: 407–409.

20 Lu FY, Wang P. Comparison of extraction process of *Hazelnut* by flash extraction and reflow method. Shaanxi Journal of Traditional Chinese Medicine. 2013; 34(4): 473–475.

21 Xie X, Li Z, Huang WC, Ye HX, Dai L, Liu XQ. Smashing tissue extraction technology optimization for polysaccharides from stems of *Acanthopanax gracilistylus* and their *in vitro* immunological activities. Chinese Traditional and Herbal Drugs. 2013; 44(20): 2859–2863.

22 Liu Y, Su ZT, Chen L, Yang L, Yang S. Study on the optimal homogenated extraction of diyu tannin by the box-behnken design. West China Journal of Pharmaceutical Sciences. 2012; 27(3):304–306.

23 Zeng ZP, Qian S, Wang H, Peng B, Han XY, Li P. Optimization of smashing tissue extraction technology of total Naphthaquinones in *Arnebia euchroma*. Chinese Journal of Experimental Traditional Medical Formulae. 2013; 19 (15): 29–31.

24 Lv DS, Meng XD, Zhang XJ. Technology optimization for extracting essential oil from *Flos Lonicera* by homogenate extraction and its application in cigarette. Chinese Agricultural Science Bulletin. 2011; 27(5): 483–488.

25 Sathishkumar P, Gu FL, Zhan QQ, Palvannan T, Yusoff ARM. Flavonoids mediated ‘Green’ nanomaterials: a novel nanomedicine system to treat various diseases-current trends and future perspective. Materials letters. 2018; 210: 26–30. https://doi.org/10.1016/j.matlet.2017.08.078

26 Tang WZ, Wang YA, Gao TY, Wang XJ, Zhao YX. Identification of C-geranylated flavonoids from Paulownia catalpifolia Gong Tong fruits by HPLC-DAD-ESI-MS/MS and their anti-aging effects on 2BS cells induced by H2O2. Chinese Journal of Natural Medicines. 2017; 15(5): 384–391. https://doi.org/10.1016/S1875-5364(17)30059-6 PMID: 28558874

27 Singh N, Rajini PS. Free radical scavenging activity of an aqueous extract of potato peel. Food Chemistry. 2004; 85(4), 611–616. https://doi.org/10.1016/j.foodchem.2003.07.003

28 Ahmed SI, Hayat MQ, Tahir M, Mansoor Q, Ismail M, Keck K, et al. Pharmacologically active flavonoids from the anticancer, antioxidant and antimicrobial extracts of Cassia angustifolia Vahl. BMC Complementary and Alternative Medicine. 2016; 16(1): 460. https://doi.org/10.1186/s12906-0161443-z PMID: 27835979

29 Pandey MM, Khatoon S, Rastogi S, Rawat AKS. Determination of flavonoids, polyphenols and antioxidant activity of Tephrosia purpurea: a seasonal study. J Integr Med. 2016; 14(6): 447–455. https://doi.org/10.1016/S2095-4964(16)60276-5 PMID: 27854196

30 Hou WC, Zhang W, Chen GD, Luo YP. Optimization of extraction conditions for maximal phenolic, flavonoid and antioxidant activity from melaleuca bracteata Leaves using the response surface methodology. PLoS One. 2016; 11(9):e0162139. https://doi.org/10.1371/journal.pone.0162139 PMID: 27611576

31 Wang YQ, Gao YJ, Ding H, Liu SJ, Han X, Gui JZ, et al. Subcritical ethanol extraction of flavonoids from moringa oleifera leaf and evaluation of antioxidant activity. Food chemistry. 2017; 218:152–158. https://doi.org/10.1016/j.foodchem.2016.09.058 PMID: 27719892

32 Deng JJ, Liu QQ, Zhang Q, Zhang C, Liu D, Fan D, et al. Comparative study on composition, physicochemical and antioxidant characteristics of different varieties of kiwifruit seed oil in China. Food chemistry. 2018; 264: 411–418. https://doi.org/10.1016/j.foodchem.2018.05.063 PMID: 29853395

33 Chung JH, Kong JN, Choi HE, Kong KH. Antioxidant, anti-inflammatory, and anti-allergic activities of the sweettasting protein brazzein. Food chemistry. 2018; 267: 163–169. https://doi.org/10.1016/j.foodchem.2017.06.084 PMID: 29934152

34 Guil-Guerrero JL, Martínez-Guirado C, Mar Rebolloso-Fuentes M, Carrique-Pérez A. Nutrient composition and antioxidant activity of 10 pepper (Capsicum annuum) varieties. European Food Research and Technology. 2006; 224(1): 1–9.

35 Song H, Zhang Q, Zhang Z, Wang J. In vitro antioxidant activity of polysaccharides extracted from Bryopsis plumosa. Carbohydrate Polymers.2010; 80: 1057–1061. https://doi.org/10.1016/j.carbpol.2010.01.024

36 Wang QZ, Yuan TF, Liu LL, Wang H, Guo J. Optimization of flash extraction of total flavonoids from leaves of podocarpus macrophyllus by response surface method and their antioxidant activity. Fine Chemicals. 2018; 35(1): 65–71.

37 Huang DN, Zhou XL, Si JZ, Gong XM, Wang S. Studies on cellulase-ultrasonic assisted extraction technology for flavonoids from Illicium verum residues. Chemistry Central Journal. 2016; 10: 56.

38 Li JJ, Zhang J, Wang M. Extraction of flavonoids from the flowers of *Abelmoschus manihot L.* medic by modified supercritical CO2 extraction and determination of antioxidant and anti-adipogenic activity. Molecules. 2016; 21(7): 810. https://doi.org/10.3390/molecules21070810 PMID: 27347916

39 Xie XJ, Zhu D, Zhang W, Huai WB, Wang K, Huang XW, et al. Microwave-assisted aqueous two-phase extraction coupled with high performance liquid chromatography for simultaneous extraction and determination of four flavonoids in *Crotalaria sessiliflora L*.. Industrial crops and products. 2017; 95: 632–642. https://doi.org/10.1016/j.indcrop.2016.11.032

40 Yedhu Krishnan R, Rajan KS. Microwave assisted extraction of flavonoids from Terminalia bellerica: Study of kinetics and thermodynamics. Separation and Purification Technology. 2016; 157:169–178. https://doi.org/10.1016/j.seppur.2015.11.035

41 Chen ZJ, Aimaiti AMGL, Dai YQ, Yuan XY, Wu YM, Ni YY. Response surface methodology as an approach to optimization of flash-extraction of flavonoids from rose fruit. Science and technology of food industry. 2011; 32(12): 387–390.

42 Ke L, Chen HY. Enzymatic-Assisted Microwave Extraction of Total Flavonoids from Bud of Chrysanthemum indicum L. and Evaluation of Biological Activities. International Journal of Food Engineering. 2016; 12(6): 607–613. https://doi.org/10.1515/ijfe-2015-0037

43 Tomaz I, Maslov L, Stupić D, Preiner D, Ašperger D, Karoglan Kontić J. Multi-response optimisation of ultrasound-assisted extraction for recovery of flavonoids from red grape skins using response surface methodology. Phytochemical analysis. 2016; 27(1): 13–22. https://doi.org/10.1002/pca.2582 PMID: 26251189

44 Sheng ZL, Wan PF, Dong CL, Li YH. Optimization of total flavonoids content extracted from *Flos Populi* using response surface methodology. Industrial crops and products. 2013; 43: 778–786. https://doi.org/10.1016/j.indcrop.2012.08.020

45 Liu YQ, Wang HW, Cai X. Optimization of the extraction of total flavonoids from Scutellaria baicalensis Georgi using the response surface methodology. J Food Sci Technol. 2014; 52(4): 2336–2343. https://doi.org/10.1007/s13197-014-1275-0 PMID: 25829617

46 Bagheri H, Manap MYBA, Solati Z. Response surface methodology applied to supercritical carbon dioxide extraction of *Piper nigrum L.* essential oil. LWT - Food Science and Technology. 2014; 57(1): 149–155. https://doi.org/10.1016/j.lwt.2014.01.015

47 Nayik GA, Dar BN, Nanda V. Optimization of the process parameters to establish the quality attributes of DPPH radical scavenging activity, total phenolic content, and total flavonoid content of apple honey using response surface methodology. International journal of food properties. 2015; 19(8): 1738–1748. https://doi.org/10.1080/10942912.2015.1107733

48 Jing CL, Dong XF, Tong JM. Optimization of ultrasonic-assisted extraction of flavonoid compounds and antioxidants from Alfalfa using response surface method. Molecules. 2015; 20(9): 15550–15571. https://doi.org/10.3390/molecules200915550 PMID: 26343617

49 Xing LJ, Liu R, Gao XG, Zheng JX, Wang C, Zhou GH, et al. The proteomics homology of antioxidant peptides extracted from dry-cured xuanwei and Jinhua ham. Food Chemistry. 2018; 266: 420–426. https://doi.org/10.1016/j.foodchem.2006.07.013

50 Salem AZM, Kholif AE, Elghandour MMY, Buendía G, Mariezcurrena MD, Hernandez SR, et al. Influence of oral administration of *Salix Babylonica* extract on milk production and composition in dairy cows. Italian journal of animal science. 2014; 13(1): 2978. https://doi.org/10.4081/ijas.2014.2978

51 Salem AZM, Olivares M, López S, González-Ronquillo M, Rojo R, Camacho LM, et al. Effect of natural extracts of *Salix babylonica* and *Leucaena leucocephala* on nutrient digestibility and growth performance of lambs. Animal Feed Science and Technology. 2011; 170(1-2): 27-34. https://doi.org/10.1016/j.anifeedsci.2011.08.002

52 Salem AZM, Robinson PH, El-Adawy MM, Hassan AA, *In vitro* fermentation and microbial protein synthesis of some browse tree leaves with or without addition of polyethylene glycol. Animal Feed Science and Technology. 2007; 138(3-4): 318–330. https://doi.org/10.1016/j.anifeedsci.2006.11.026

53 Salem AZM, Salem MZM, González-Ronquillo M, Camacho LM, Cipriano M. Major chemical constituents of Leucaena leucocephala and *Salix babylonica* leaf extracts. Journal of Tropical Agriculture. 2011; 49:95.

54 Zheng YN, Zhang J, Han LK, Sekiya K, Kimura Y, Okuda H. Effects of compounds in leaves of Salix matsudana on arachidonic acid metabolism. Yakugaku Zasshi. 2005; 125(12): 1005–1008. https://doi.org/10.1248/yakushi.125.1005 PMID:16327246

55 Li X, Liu Z, Zhang XF, Wang LJ, Zheng YN, Yuan CC, et al. Isolation and characterization of phenolic compounds from the leaves of *Salix matsudana.* Molecules. 2008; 13(8): 1530–1537. https://doi.org/10.3390/molecules13081530 PMID: 18794770

56 Kaufeld AM, Pertz HH, Kolodziej H. A chemically defined 2, 3-trans procyanidin fraction from willow bark causes redox-sensitive endothelium-dependent relaxation in porcine coronary arteries. Journal of Natural Products. 2014; 77(7): 1607–1614. https://doi.org/10.1021/np500177u PMID: 24957134

57 Sharma S, Sahu D, Das HR, Sharma D. Amelioration of collagen-induced arthritis by *Salix nigra* bark extract via suppression of pro-inflammatory cytokines and oxidative stress. Food Chemistry. 2011; 49(12): 3395–3406 https://doi.org/10.1016/j.fct.2011.08.013 PMID: 21983485

58 Freischmidt A, Jurgenliemk G, Kraus B, Okpanyi SN, Muller J, Kelber O, et al. Contribution of flavonoids and catechol to the reduction of ICAM-1 expression in endothelial cells by a standardised Willow bark extract. Phytomedicine. 2012; 19(3-4): 245–252. https://doi.org/10.1016/j.phymed.2011.08.065 PMID: 21982436

59 Enayat S, Banerjee S. The ethanolic extract of bark from *Salix aegyptiaca L*. inhibits the metastatic potential and epithelial to mesenchymal transition of colon cancer cell lines. Nutrition and Cancer-AN International Journal. 2014; 66(6): 999–1008. https://doi.org/10.1080/01635581.2014.936949 PMID: 25175673

60 Kong CS, Kim KH, Choi JS, Kim JE, Park C, Jeong JW. Salicin, an Extract from White Willow Bark, Inhibits Angiogenesis by Blocking the ROS-ERK Pathways. Phytother. Res. 2014; 28(8): 1246. https://doi.org/10.1002/ptr.5126 PMID: 24535656

61 Ilnicka A, Roszek K, Olejniczak A, Komoszynski M, Lukaszewicz JP. Biologically active constituents from *Salix* viminalis bio-oil and their protective activity against hydrogen peroxide-induced oxidative stress in Chinese hamster ovary cells. Applied biochemistry and biotechnology. 2014; 174(6): 2153–2161. https://doi.org/10.1007/s12010-014-1171-0 PMID: 25172057

62 Dziki UG, Sugier D, Dziki D, Sugier P. Bioaccessibility In Vitro of Nutraceuticals from Bark of Selected Salix Species. The Scientific World Journal. 2014; 782763. http://dx.doi.org/10.1155/2014/782763 PMID: 24696660

63 Yang H, Lee SH, Sung SH, Kim J, Kim YC. Neuroprotective compounds from Salix pseudo-lasiogyne twigs and their anti-amnesic effects on scopolamine-induced memory deficit in mice. Planta Medica. 2013; 79(1): 78–82. http://dx.doi.org/10.1055/s-0032-1327949 PMID: 23154841

64 Lee M, Lee SH, Kang J, Yang H, Jeong EJ, Kim HP. Salicortin-derivatives from Salix pseudo-lasiogyne twigs inhibit adipogenesis in 3T3-L1 cells via modulation of C/EBP-αandSREBP1-c dependent pathway. Molecules. 2013; 18(9): 10484–10496. http://dx.doi.org/10.3390/molecules180910484 PMID: 23999723

65 Liu KY, Zhou B, Liu HJ. Studies on the extraction technology and determination of total flavonoids in *Salix babylonica L.* Journal of Anhui Agricultural Sciences. 2007; 35(31): 9813, 9823.

66 Shen R. Progress in the application of flash extraction in traditional Chinese medicine. Journal of Chinese Medicinal Materials. 2015; 38(7): 1540–1542.

67 Huang YY, Tan RD. Research progress of flash extraction of active ingredients from Chinese herbal medicine. Chinese Journal of Hospital Pharmacy. 2015; 35(1): 83–87.

68 Duan MH, Ma JF, Zhang C, Ge FH. Research progress on pretreatment methods of active ingredient samples of traditional Chinese medicine. Journal of Chinese Medicinal Materials, 2017; 40(6): 1495–1498.

69 Ren YF, Yi H, Gao J, Yang H. Study on Application of Smashing Tissue Extraction in Extracting Fat-Soluble Compounds of Salvia Miltiorrhiza by Microemulsion. Chinese Journal of Experimental Traditional Medical Formulae. 2011; 17(2): 15–20.

70 Pham-Huy LA, He H, Pham-Huy C. Free radicals, antioxidants in disease and health. International journal of biological sciences. 2008; 4(2): 89–96. PMID: 23675073

71 Xie J, Zhou PP, Zhu XY, Liu XJ, Chen, RE, Wang P. Study on extraction of bamboo leaves flavonoids by homogenate extraction technique and its antioxidant activity. Food science and technology. 2010; 35(5): 194–198.

